# A human pluripotent stem cell tri-culture platform to elucidate microglial regulation of retinal ganglion cells in neuroinflammation

**DOI:** 10.1101/2025.07.01.662573

**Authors:** Jade Harkin, Cátia Gomes, Kaylee D. Tutrow, Reham Afify, Shruti V. Patil, Kiersten H. Peña, Aaron Baker, Sailee S. Lavekar, Kang-Chieh Huang, Jason S. Meyer

## Abstract

Optic neuropathies, including glaucoma, are characterized by the progressive degeneration of retinal ganglion cells (RGCs), ultimately leading to irreversible vision loss. Increasing evidence implicates microglia, the resident immune cells of the central nervous system, as key modulators of RGC health and disease progression. However, the precise mechanisms by which microglia influence RGCs remain poorly understood, particularly in the human context. In this study, we established human pluripotent stem cell (hPSC)-derived co-culture systems incorporating microglia, astrocytes, and RGCs to explore how microglia shape RGC growth and maturation under physiological conditions. We first examined the impact of homeostatic microglia on RGCs in both co-culture and tri-culture systems, revealing distinct influences of cell types in co-culture compared to when they were grown individually. We then modeled inflammatory states by activating microglia with lipopolysaccharide (LPS) and evaluated their effects on RGCs both directly and in the context of astrocyte co-culture. This stepwise, reductionist approach enabled us to dissect the cellular interactions driving RGC vulnerability in inflammatory conditions relevant to optic neuropathies. Our findings provide new insight into the complex neuroimmune landscape that underlies RGC degeneration and identify key pathways that may serve as therapeutic targets across a range of optic nerve diseases.

## INTRODUCTION

Microglia are the principal immune cells of the central nervous system (CNS), where they play critical roles in maintaining homeostasis, surveying the local environment, and responding to injury or disease (1, 2). Unlike peripheral macrophages, microglia are uniquely adapted to the CNS environment and contribute to neural development and maintenance by regulating synapse formation, pruning, and neuronal survival (3, 4). The retina, as a specialized extension of the CNS, is composed of multiple neural and glial cell types and is connected to the brain via the optic nerve, formed by the axons of retinal ganglion cells (RGCs) (5, 6). Within the retina and optic nerve, microglia reside predominantly in the plexiform layers and the optic nerve head, where they exhibit a ramified morphology under healthy conditions (7, 8). During development, microglia work in concert with astrocytes to shape retinal circuits by promoting RGC maturation and eliminating superfluous synapses. Both glial populations play essential roles in supporting the structural and functional integrity of the retina (9, 10). However, in response to injury, stress, or disease, microglia transition to an activated pro-inflammatory state, characterized by altered morphology, upregulation of inflammatory signaling pathways, and secretion of cytokines and neurotoxic factors (7, 11). While these responses may serve protective or reparative functions in acute contexts, sustained microglial activation contributes to chronic neuroinflammation and RGC degeneration, with this inflammatory response enhanced by microglial activation of neighboring astrocytes (7, 12). Understanding how microglia transition between supportive and pathogenic states, as well as how they modulate the growth or degeneration of RGCs, has been a key point of emphasis to better understand their roles in both normal retinal development and the pathogenesis of optic neuropathies.

Much of our current understanding of how microglia modulate RGCs and interact with astrocytes has been derived from studies in rodent models. These models have been instrumental in revealing how microglia influence RGC development, synaptic refinement, and responses to injury, as well as how microglia-astrocyte signaling contributes to both neuroprotection and neurodegeneration. However, significant anatomical and molecular differences exist between rodent and human retinas and optic nerves (13, 14), limiting the translational relevance of these findings. Post-mortem human tissue has provided important insights into microglial activation in disease (15, 16), but such tissue typically reflects end-stage degeneration and does not permit dynamic or causal experimentation. Consequently, alternative approaches are needed to model human specific glial-neuronal interactions under both physiological and pathological conditions. Human pluripotent stem cells (hPSCs) offer a powerful platform to address this gap, enabling the generation of microglia, astrocytes, and RGCs from the same genetic background and the ability to study their interactions in a highly controlled and experimentally tractable system.

Thus, to address this shortcoming and advance our understanding of how human microglia modulate developmental and disease states of RGCs, we have developed novel co-culture models of hPSC-derived microglia, astrocytes and RGCs to explore the cellular crosstalk between these cell types. Microglia were differentiated from hPSCs and when grown in co-culture with RGCs, they resulted in greater neuronal growth and functional maturation. Similarly, when astrocytes were added to generate a tri-culture system, we found that microglia further regulated astrocyte growth, which in turn contributed to RGC development and maturation. Conversely, we were successfully able to induce a pro-inflammatory profile in hPSC-derived microglia, which resulted in significant degenerative phenotypes within RGCs, as well as the concurrent activation of astrocytes in tri-cultures. Furthermore, we explored the interactions of all three cell types in tri-cultures in developmental and neuroinflammatory contexts by single cell RNA-seq analyses, identifying candidate gene networks activated in each cell type when grown in these conditions. Taken together, this study is the first study to generate a human cellular model system that allows for a more sophisticated investigation of cellular interactions between human microglia, astrocytes and RGCs that would otherwise be inaccessible for investigation, providing novel opportunities for the development of new experimental approaches to understand mechanisms of neuroinflammation in the retina and optic nerve.

## RESULTS

### Microglia Enhance Retinal Ganglion Cell Morphological Complexity and Functional Maturation

To explore the role of microglia in the regulation of RGC development and maturation, we adopted a well-established differentiation protocol described by McQuade et al. (17), in which MGLs are derived through a stepwise process initially yielding enriched populations of hematopoietic progenitor cells, followed by their subsequent differentiation into microglia (Figure 1A). To confirm the microglial identity of resulting cells, we performed qRT-PCR analyses of several microglial-associated genes and compared these results to similar experiments run on undifferentiated hPSCs, as well as hPSC-derived neurons and astrocytes (Figure 1B). In these experiments, we found that these microglial associated transcripts were highly expressed only in those cells derived from microglia differentiation paradigms. Successful differentiation was then further confirmed via immunocytochemistry, which demonstrated robust expression of microglia-specific markers P2RY12, IBA1, and TREM2 in nearly all differentiated cells (Figure 1C–E).

**Figure 1.**
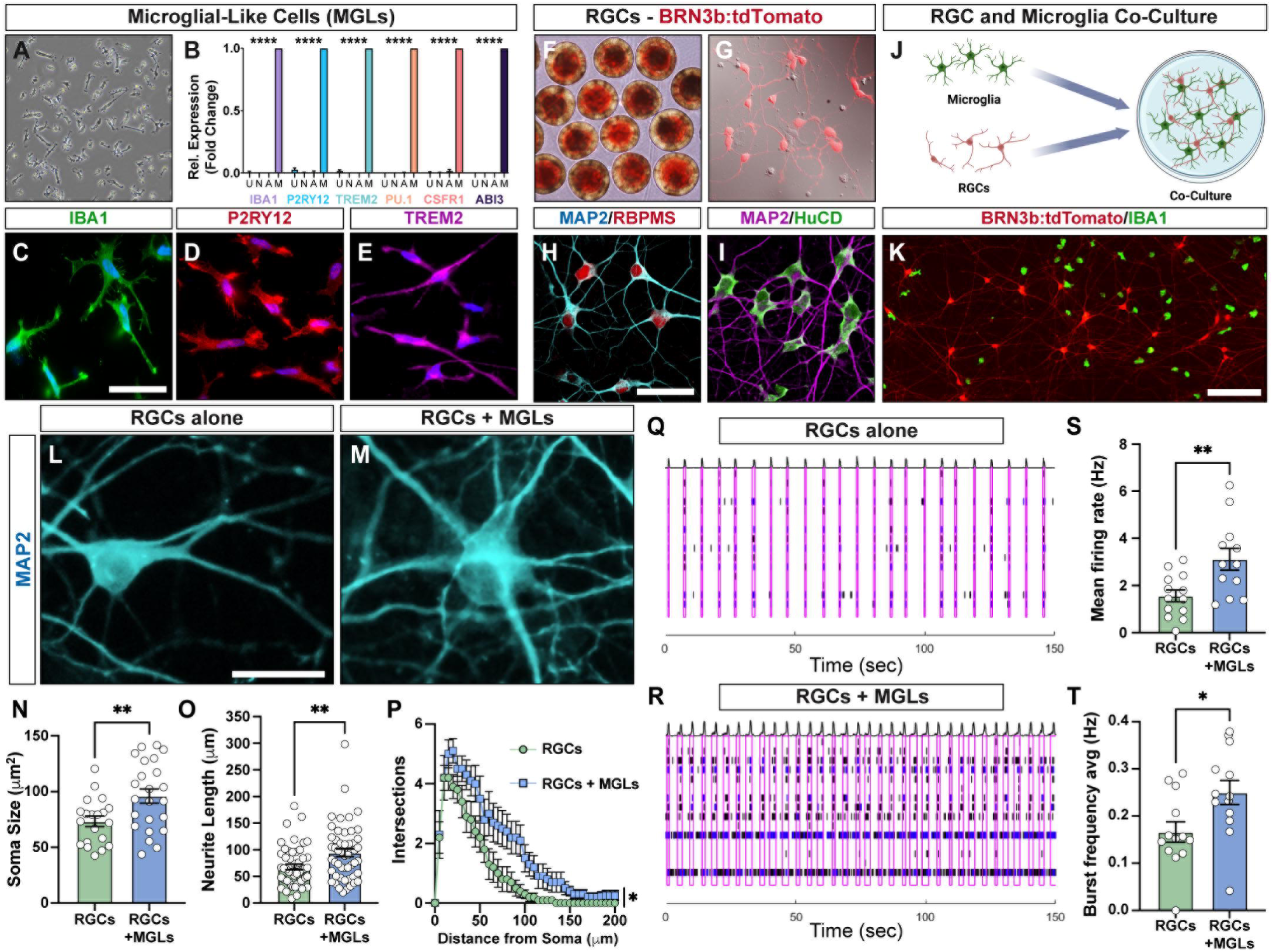
hPSC-derived microglia support RGC neurite outgrowth and function. (A) Representative brightfield image of microglia (MGLs) differentiated from hPSCs. (B) qRT-PCR analyses for P2RY12, IBA1, TREM2, PU.1, CSFR1 and ABI3 to further confirm that MGLs express microglial associated genes compared to negative controls (undifferentiated iPSCs, astrocytes and cortical neurons). (C-E) Immunostaining of microglia-like cells for microglia markers, IBA1, TREM2 and P2RY12. (F-G) Representative images of retinal organoids as well as isolated 2D cultures of BRN3b:tdTomato RGCs. (H-I) Immunostaining of hPSC-derived RGCs for characteristic markers such as MAP2, RBPMs, MAP2 and HuCD. (J) Schematic representation of hPSC-derived microglia and RGC co-cultures. (K) Immunostaining of IBA1-positive microglia and BRN3:tdTomato-RGCs in co-culture. (L-M) Representative images of RGCs co-cultures with microglia. (N-O) Quantification of RGC morphology including soma size and neurite length of RGC co-cultured with microglia or grown alone. (P) Sholl analysis of RGC neurite complexity grown in co-culture with microglia. (Q-R) Representative raster plots of RGCs when co-cultured with MGLs or grown alone showing that microglia promote an increase in RGC network bursting. Black bars represent spikes, blue bars represent bursts, and pink rectangles represent network bursting. (S-T) Quantification of RGC functional properties including mean firing rate and average burst frequency when co-cultured with microglia or grown alone. Data represent mean values ± SEM. *p < 0.05 and **p < 0.01. Scale bar equals 30 μm in C, H and L, 200 μm in K.

Concurrently, we derived RGCs from hPSCs with a Brn3b-tdTomato–Thy1.2 selectable marker, as previously described (18, 19). Initially, retinal organoids were derived following established protocols, and RGCs could be readily identified based upon the expression of the tdTomato reporter in the inner layers of retinal organoid structures (Figure 1F). Retinal organoids were then enzymatically dissociated and RGCs were purified by MACS sorting against the Thy1.2 antigen, yielding highly enriched populations of RGCs expressing a variety of RGC-associated markers (Figure 1G-I). To investigate the influence of microglia on RGCs, we then established a co-culture system using hPSC-derived RGCs and microglia (Figure 1J–K). Compared to RGCs grown alone, RGCs grown in co-culture with microglia exhibited markedly enhanced neuronal morphology, including significantly increased soma size, neurite length, and neurite complexity as assessed by Sholl analysis (Figure 1L–P). To determine if the co-culture of RGCs with microglia led to functional changes in RGCs, we assessed these cultures by multielectrode array (MEA), which demonstrated enhanced electrophysiological activity of RGCs in co-culture with microglia, including an increased mean firing rate as well as burst frequency (Figure 1Q–T).

### Establishment of a Tri-culture Model to Investigate Cellular Interactions Between Retinal Ganglion Cells, Astrocytes, and Microglia

Building on our findings that microglia enhance the maturation of stem cell-derived RGCs in co-culture, we next examined how microglia influence astrocytes and how these interactions shape the collective behavior of all three cell types. To explore this, we initially established co-cultures between hPSC-derived astrocytes and microglia (Figure 2A). Notably, when co-cultured with microglia, hPSC-derived astrocytes exhibited more complex morphological features characterized primarily by increased branch length (Figure 2B-D). Given this influence of microglia over astrocytes, as well as our previous studies demonstrating the growth and maturation inducing effects of astrocytes upon RGCs (20–22), we then established a tri-culture model of RGCs, microglia, and astrocytes to investigate the interactions between these cell types and how these interactions influence cellular growth and maturation (Figure 2E–F). We compared each cell type grown in monoculture to those grown in the tri-culture system under standardized media conditions, where we observed that cells in the tri-culture system exhibited features consistent with enhanced morphological maturation. RGCs showed significantly increased neurite length and branching complexity (Figure 2G–J), astrocytes exhibited more elaborate morphologies with an increased number of processes which were also longer (Figure 2K–N), and microglia displayed more highly ramified morphological features with increased process length and endpoint number (Figure 2O–R). These findings suggest that cell–cell communication in tri-culture promoted more physiologically relevant maturation compared to when these cells were grown in monoculture alone, supporting the tri-culture model as a more robust in vitro system to study dynamic interactions between RGCs and glia.

**Figure 2.**
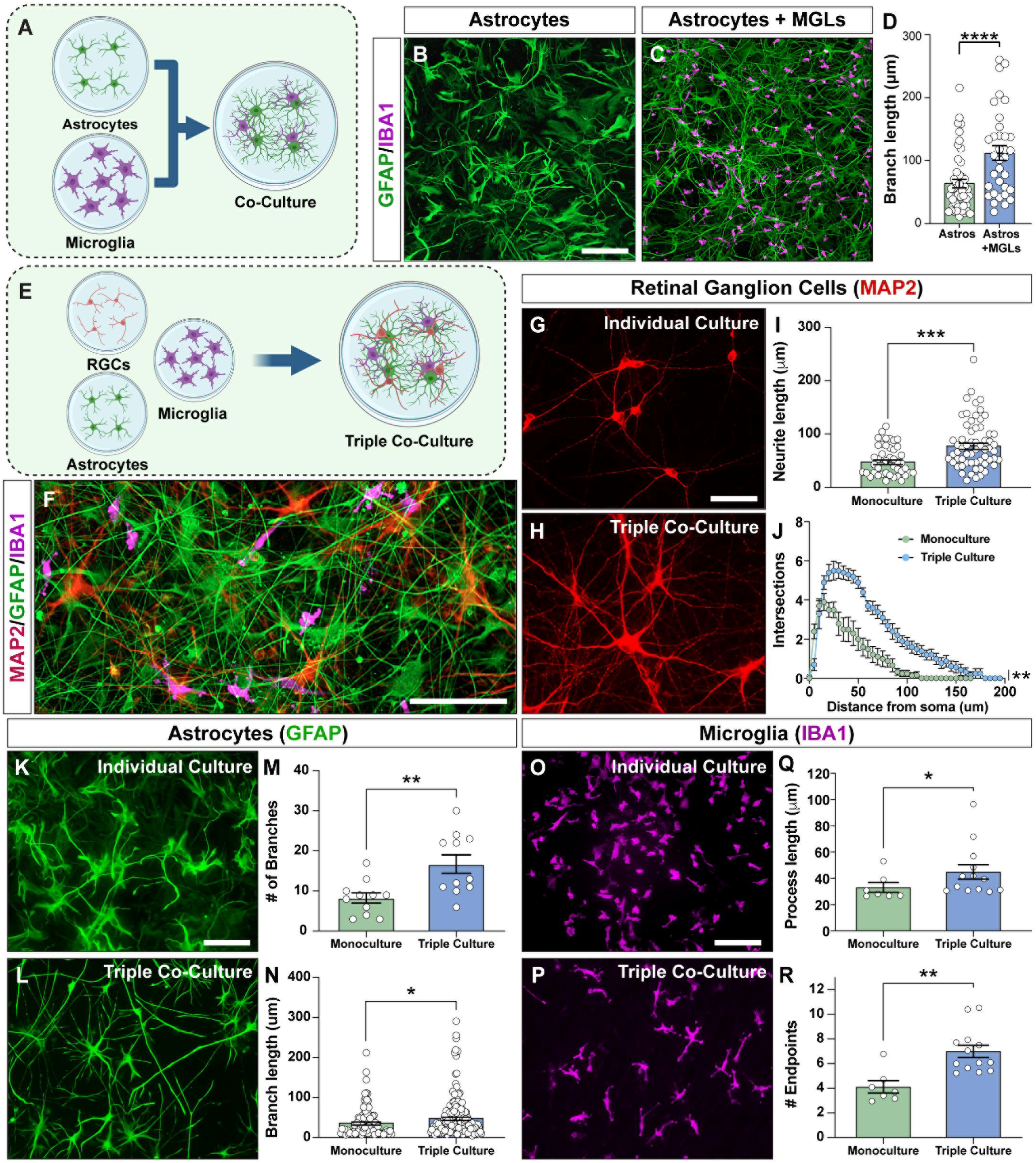
Development of a tri-culture system to study interactions among RGCs, astrocytes and microglia. (A) Schematic of hPSC-derived astrocytes and MGL co-cultures. (B-C) Representative images of astrocytes (GFAP) alone or in co-culture with MGLs (IBA1). (D) Morphological analysis of astrocytes showed that MGLs enhance branch length in astrocytes. (E) Schematic of hPSC-derived RGC, astrocyte and MGL triple cultures. (F) Representative images of RGCs (MAP2) astrocytes (GFAP) and MGLs (IBA1) grown in triple-culture. (G-H) Representative images of RGCs (MAP2) grown alone or in triple-culture with astrocytes and MGLs showing increased complexity when in contact with glial cells. (I-J) Quantification of RGC morphology including neurite length and Sholl analysis, in monoculture or in triple culture with astrocytes and MGLs. (K-L) Representative images of astrocytes (GFAP) grown in monoculture or in triple-culture with RGCs and MGLs. (M-N) Analysis of astrocyte morphology, including number of branches and branch length, when grown alone or in triple-cultures. (O-P) Representative images of MGLs (IBA1) in monoculture or in triple-culture with astrocytes and RGCs. (Q-R) Morphological analysis of MGLs, including process length and number of endpoints when grown alone or in triple-cultures. Data represent mean values ± SEM. *p < 0.05, **p < 0.01, ***p < 0.001 and ****p < 0.0001. Scale bar equals 50 μm in B, G, K, and O, and 100 μm in F.

### Single Cell RNA-seq profiling of RGCs, microglia, and astrocytes to identify transcriptional correlates of cellular maturation in tri-culture

To further investigate how interactions between RGCs and glial cells influence gene expression profiles, we conducted single-cell RNA sequencing analyses on RGCs, astrocytes, and microglia each grown individually in monoculture, and compared them to tri-cultures containing all three cell types (Figure 3A). This approach enabled us to examine how the tri-culture environment impacted gene expression and cellular maturation of each specific cell type. In these analyses, populations of RGCs, astrocytes, and microglia were identified as distinct populations (Figure 3B), and we then explored differentially expressed genes in each cell type between monoculture and tri-culture conditions, with many transcripts upregulated in tri-culture, while others were upregulated in individual cells in monoculture (Figure 3C). Among RGCs which were identified based upon the expression of transcripts including POU4F2, SNCG, POU6F2, and SIX6 (23, 24), we observed numerous transcriptional changes (Figure 3D-E). Interestingly, a range of RGC maturation stages seemed to be evident in these analyses, with some more immature cells retaining SIX6 expression, while more differentiated RGCs expressed POU4F2. This spectrum of RGC maturation played out in comparisons between RGCs grown in monoculture compared to those grown in tri-culture, where more retinal development-associated genes such as VSX2, RAX, ATOH7 and SIX6 (25–27) were upregulated in RGCs grown in monoculture, whereas RGCs grown in tri-culture expressed more maturation-associated transcripts, including NEFL, RBPMS, and SYN1 (28, 29). Similarly, genes associated with axonal growth (PLXNA4, SHTN1, RHEB, NTRK2) and neuronal excitability (CANA2D3, GABRA2) were upregulated in RGCs grown in tri-cultures, further supporting the maturation-inducing phenotypes of RGCs induced by glia.

**Figure 3.**
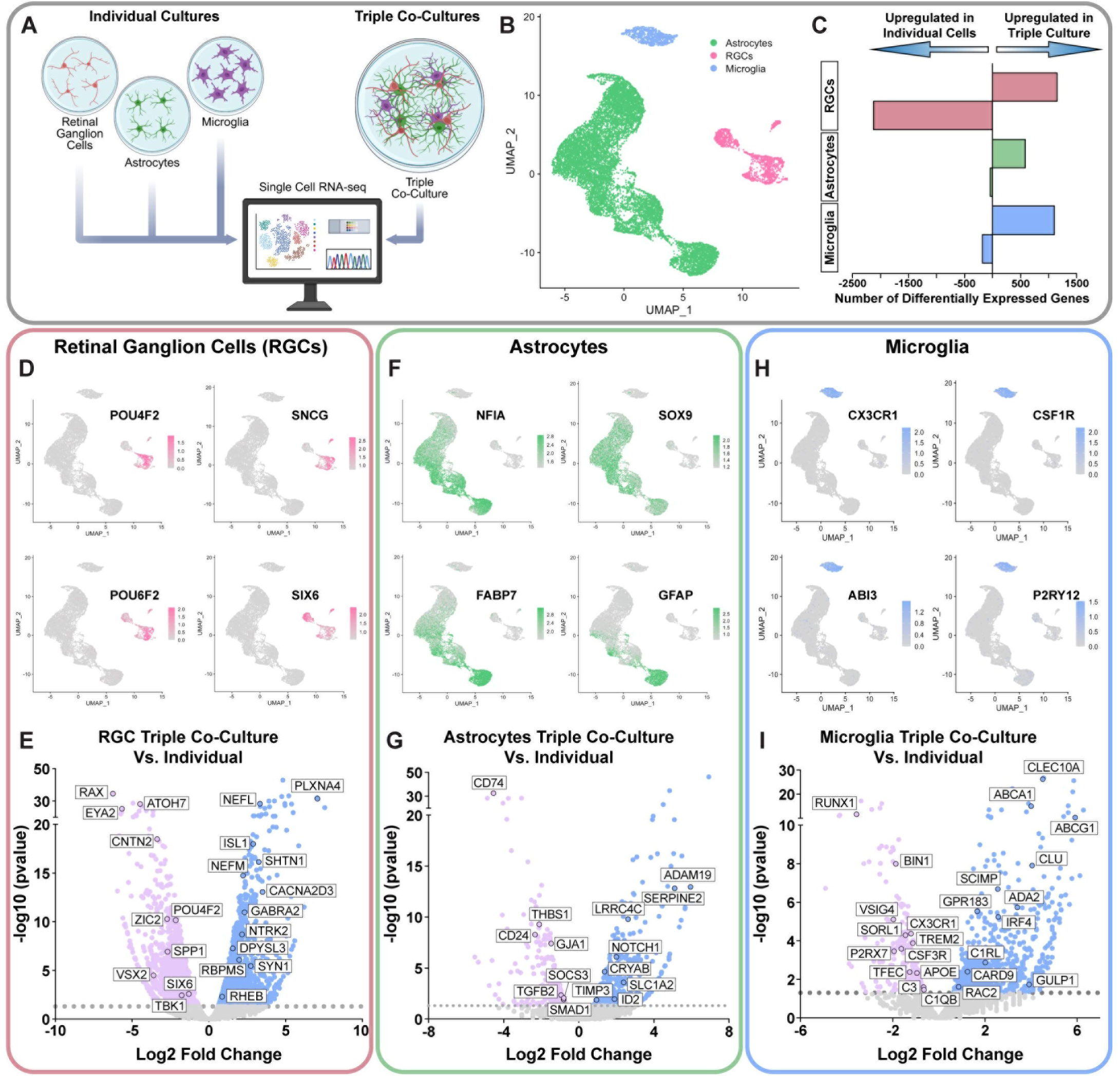
scRNA-seq of RGCs, MGLs and astrocytes to uncover transcriptional signatures associated with cellular maturation in a tri-culture system. (A) Schematic of cell culture systems used for scRNA-seq including RGCs, astrocytes and MGLs grown individually in monoculture, as well as tri-cultures containing all three cell types. (B) UMAP projection of scRNA-seq representing RGC, astrocyte and MGL population colored in pink, green and blue, respectively. (C) Representation of the number of differentially expressed genes for each cell type grown in monoculture or in tri-culture. (D) UMAPs representing RGC populations identified by characteristic genes including POU4F2, SNCG, POU6F2 and SIX6. (E) Volcano plot exhibiting differentially expressed genes in RGC monocultures compared to RGCs grown in triple co-cultures with astrocytes and MGLs. Significantly upregulated genes are shown in blue, while significantly downregulated genes are shown in pink. (F) UMAPs representing astrocyte populations identified by characteristic genes such as NFIA, SOX9, FABP7 and GFAP. (G) Volcano plot exhibiting differentially expressed genes in astrocyte monocultures compared to astrocytes grown in triple co-cultures with RGCs and MGLs. (H) UMAPs representing MGL populations identified by characteristic genes including CX3CR1, CSF1R, ABI3 and P2RY12. (I) Volcano plot exhibiting differentially expressed genes in MGL monocultures compared to MGLs grown in triple co-cultures with RGCs and astrocytes.

Concurrently, we also explored how the transcriptional profile of glia was modulated in tri-culture compared to growth of each cell type in monoculture. Similar to the developmental spectrum identified in RGCs, we similarly found a range of maturity among astrocytes, with nearly all astrocytes expressing pan-astrocyte genes including NFIA and SOX9, whereas more mature astrocytes expressed FABP7 and GFAP (Figure 3F-G). Among astrocytes grown in tri-culture, we identified numerous genes associated with modulation of the extracellular matrix (SERPINE2, ADAM19, TIMP3) as well as astrocyte specification (NOTCH1, ID2). Conversely, astrocytes grown in monoculture expressed more genes associated with astrocyte activation (TGFb2, SOCS3, CD74). We did not observe a change in expression of genes more classically associated with an astrocytic activation profile such as complement component C3 (12, 20), suggesting that these other activation-associated genes more likely imply that astrocytes grown in tri-culture are simply more homeostatic than those grown in monoculture.

Furthermore, we assessed how microglia grown in a tri-culture system modulated microglial signatures compared to microglia grown in mono-culture (Figure 3H-I). Microglia were identified in these analyses based upon the expression of canonical markers including CX3CR1, CSF1R, ABI3, and P2RY12 (Figure 3H). When we compared microglia grown in tri-culture to those grown in mono-culture, we found that microglia grown in tri-culture appeared more mature exhibiting higher expression of genes associated with microglial function, including those associated with lipid metabolism (ABCA1, CLU, ABCG2), innate immune signaling (SCIMP, ADA2, RAC2, CARD9), and phagocytosis (GULP1, CLEC10A). Conversely, microglia grown in monoculture expressed higher levels of transcripts for genes associated with neurodegenerative diseases, including BIN1, TREM2, SORL1, and APOE, suggesting that microglia grown in monoculture may be more primed for disease-related phenotypes or conversely, that microglia grown in tri-culture with astrocytes and RGCs maintained a more homeostatic profile.

### Stimulating a pro-inflammatory signature in microglia to explore neuroinflammatory phenotypes

While microglia are known to play essential roles in the maintenance of a homeostatic state in the central nervous system, they also play pivotal roles in neurodegenerative states including glaucoma, where they adopt an inflammatory profile and contribute to the neurodegeneration of RGCs (30, 31). To explore how activated microglia may adversely affect RGCs, we initially sought to replicate this pro-inflammatory activated state in hPSC-derived microglia by treatment with LPS, at which point they exhibited increased MHC-II expression as a characteristic of an activated microglial state (Figure 4A-D) (32, 33). Additionally, qRT-PCR analyses demonstrated that LPS-activated MGLs also increased expression of the MHC-II transporter, CD74 (Figure 4E), further supporting the ability to induce this activated microglial state (34). As activated microglia increase the secretion of pro-inflammatory cytokines (7, 12, 35), next we then wanted to confirm whether treatment with LPS led to similar increases in hPSC-derived microglia. After 24 hours of LPS treatment, we collected conditioned media from both control and LPS-activated microglia and analyzed secreted factors released from these cells using the Meso Scale Discovery (MSD) V-Plex proinflammatory panel (Figure 4F-G), with results demonstrating that LPS-activated MGLs released increased concentrations of many pro-inflammatory cytokines compared to control MGLs. Further, to assess the phagocytic activity of both control and LPS-activated MGLs, cells were first incubated with LPS for 24 hours or maintained as control, non-stimulated cultures, followed by incubation with pHrodo Zymosan-Red BioParticles to assess the phagocytic capacity of these cells (Figure 4H-M, Supplemental Movie 1). Results showed that LPS-activated microglia were more highly phagocytic at all time points when compared to control microglia (Figure 4N).

**Figure 4.**
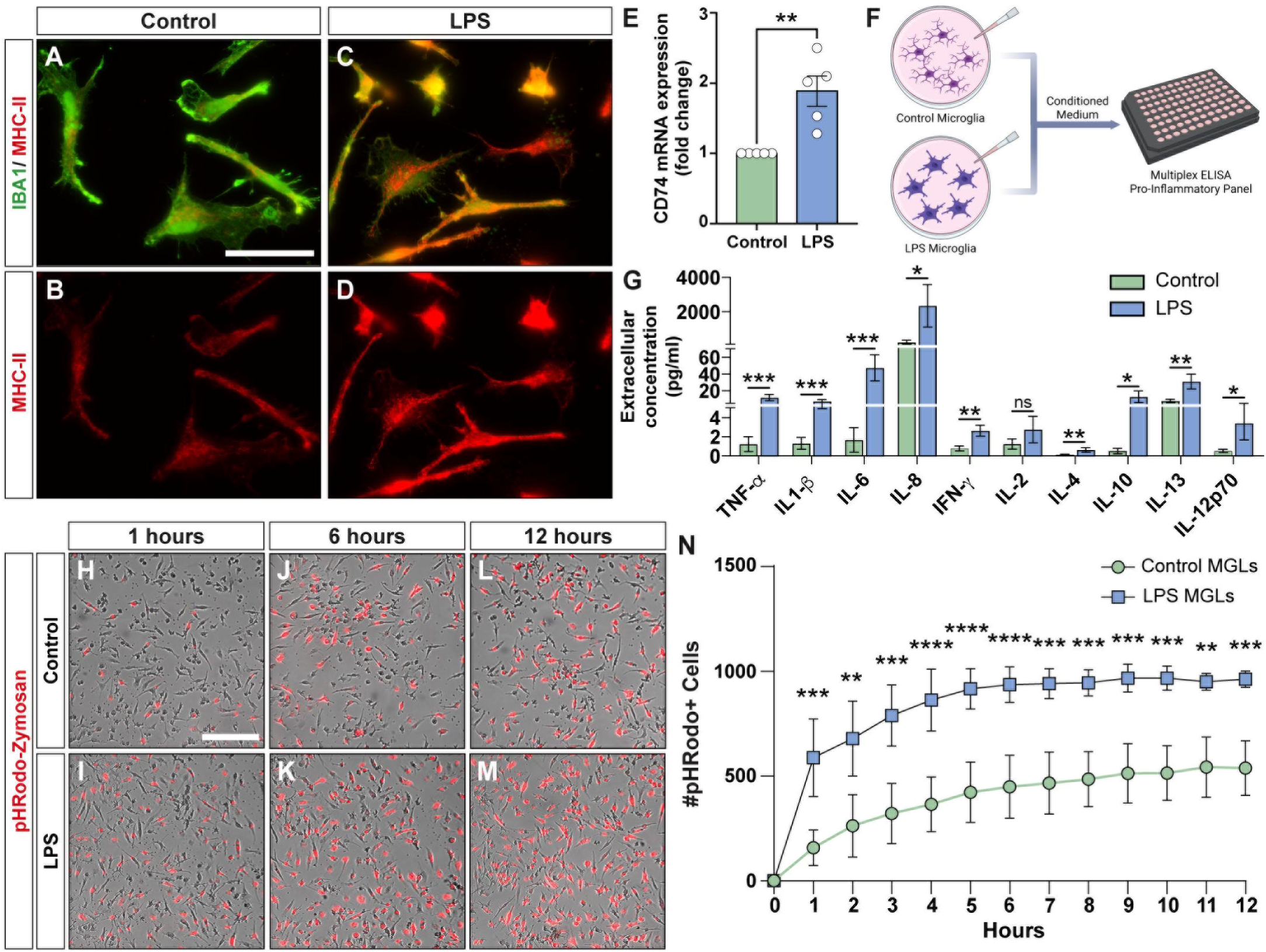
Induction of a pro-inflammatory phenotype in hPSC-derived microglia. (A-D) Immunostaining of MGLs stimulated with LPS (150 ng/ml) for 24 hours, showing that LPS-treated MGLs display increased MHC-II expression compared to control MGLs. (E) qRT-PCR analyses of CD74 (MHC-II transporter) expression in LPS-stimulated and control MGLs. (F) Schematic representing the collection of extracellular conditioned media of LPS-treated MGLs and controls and their analysis through multiplex ELISA. (G) Meso Scale Discovery (MSD) V-Plex proinflammatory panel analyzing the secretion of inflammatory cytokines in conditioned media collected from control MGLs and LPS-stimulated MGLs. (H-M) Representative time-lapse images showing pHrodo Zymosan-Red bioparticles engulfed by LPS-stimulated MGLs or respective controls, at 1 hour, 6 hours and 12 hours. (N) Quantification shows the red object count for the phagocytosis of pHrodo Zymosan-Red bioparticles in control MGLs compared to LPS MGLs over a 12-hour time period. Data represent mean values ± SEM. *p < 0.05, **p < 0.01, ***p < 0.001 and ****p < 0.0001, ns = not significant. Scale bar equals 50 μm in A and 200 μm in H.

### Pro-inflammatory microglia promote RGC neurodegenerative phenotypes

In glaucomatous damage under chronic inflammatory conditions, microglia activation contributes to RGC neurodegeneration and tissue damage (7, 30). Therefore, we initially established co-cultures to examine the effects of microglial activation on RGCs (Figure 5A-B). When LPS-stimulated microglia were co-cultured with RGCs for 3 weeks, we observed detrimental effects of LPS-activated microglia on RGC morphological features, with RGCs exhibiting reduced soma size, neurite length and number of primary neurites compared to RGCs that were co-cultured with control microglia (Figure 5C-E). Next, we wanted to determine if pro-inflammatory microglia resulted in any functional deficits in RGCs. Similar co-cultures were then established on an MEA platform to examine the functional properties of RGCs grown with either control or LPS-activated microglia for 3 weeks (Figure 5F-G). Notably, we observed that LPS-activated pro-inflammatory microglia reduced the mean firing rate of RGCs compared to controls (Figure 5H-J). Moreover, the average burst frequency and the average burst percentage of RGCs was significantly when co-cultured with LPS-stimulated pro-inflammatory microglia (Figure 5K-L), further supporting the concept that pro-inflammatory microglia can contribute to RGC neurodegeneration.

**Figure 5.**
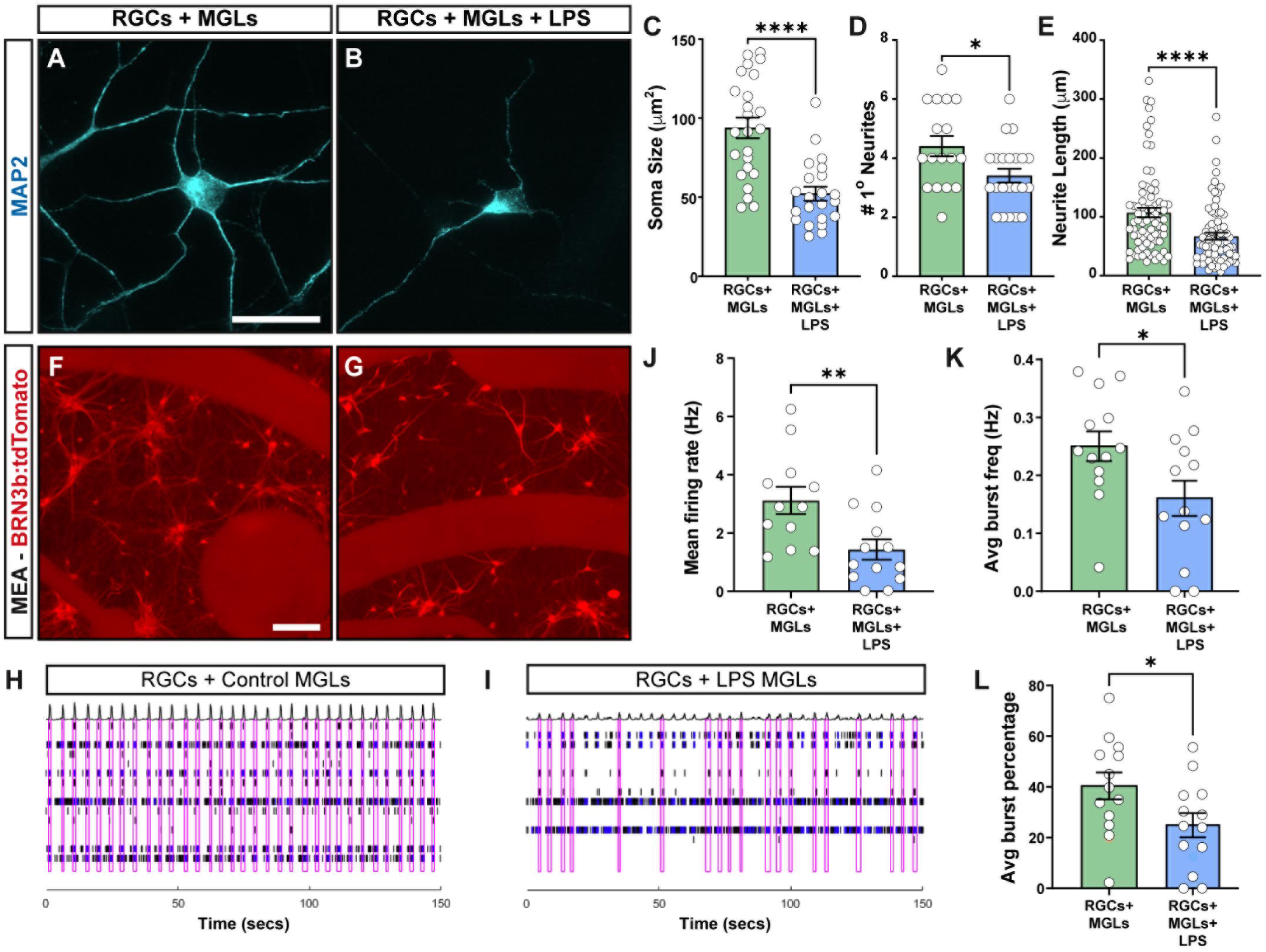
Pro-inflammatory MGLs promote RGC neurodegenerative phenotypes. (A-B) Representative images of RGCs (MAP2) co-cultured with LPS-stimulated MGLs or respective controls. (C-E) Quantification of RGC morphology including soma size, number of primary neurites and neurite length grown in co-culture with MGLs and treated with or without LPS. (F-G) Representative images of RGCs (BRN3b:tdTomato) in multielectrode array (MEA) plates in co-culture with MGLs treated with LPS or untreated controls. (H-I) Representative raster plots of RGCs when co-cultured with LPS-stimulated MGLs or respective controls, showing that LPS-treated NGLs promote a decrease in RGC network bursting. Black bars represent spikes, blue bars represent bursts, and pink rectangles represent network bursting. (J-L) Quantification of RGC functional properties including mean firing rate, average burst frequency and average burst percentage when co-cultured with LPS-treated MGLs or non-treated control. Data represent mean values ± SEM. *p < 0.05, **p < 0.01 and ***p < 0.0001. Scale bar equals 30 μm in C and 200 μm in F.

### Use of the tri-culture model to investigate interactions between RGCs, astrocytes, and microglia under neuroinflammatory conditions

It is well-established that microglia and astrocytes can work in concert to create a neuroinflammatory environment, resulting in the degeneration of neurons such as RGCs. However, these studies are often performed in animal models, which lack the ability to experimentally isolate individual cell types and assess contributions to degenerative states. Previous studies have demonstrated that neurotoxic astrocytes are induced by pro-inflammatory microglia through the secretion of factors including TNFα, IL-1α and C1q (12, 36), suggesting that the combined actions of pro-inflammatory microglia and astrocytes may create a more robustly neurotoxic environment. However, these studies have not yet been performed in human stem cell-derived microglia, astrocytes, and RGCs, particularly in a manner that is conducive to the exploration of how these cell types are each individually modulated in this environment.

To address this, we initially established co-cultures of hPSC-derived astrocytes and microglia to examine the cellular crosstalk between these cell types in control or pro-inflammatory states (Figure 6A). Initially, astrocytes were co-cultured with LPS-activated microglia for 7 days before changes in astrocyte morphological phenotypes were assessed. Results demonstrated that when directly co-cultured with LPS-activated microglia, astrocytes exhibited a more compacted, reactive-like phenotype, characterized by an increase in astrocyte soma area as well as a decrease in the individual astrocyte branch length (Figure 6B-E), compared to astrocytes co-cultured with control microglia. Importantly, control experiments of astrocytes grown in monoculture in the presence of LPS did not demonstrate any morphological changes compared to astrocytes grown alone, confirming that LPS stimulation only acted through microglia, at least in the context of these experimental outputs, and any changes in astrocytes were influenced by microglia.

**Figure 6.**
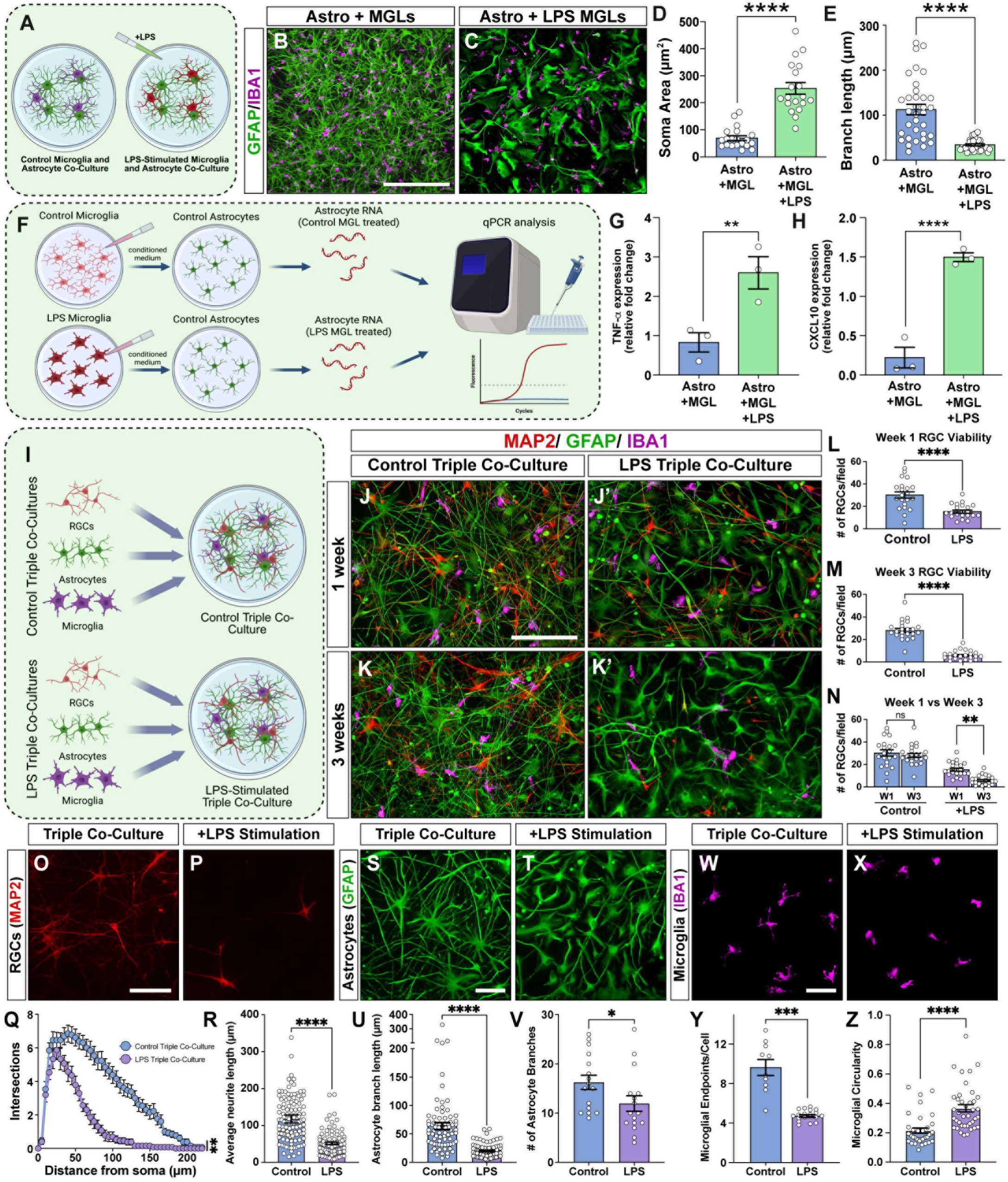
Induction of a pro-inflammatory environment in tri-culture of RGC, astrocytes and microglia. (A) Schematic of hPSC-derived astrocyte and MGL control co-cultures or stimulated with LPS. (B-C) Representative images of co-cultures containing astrocytes (GFAP) and MGLs (IBA1), treated with and without LPS. (D-E) Quantification of astrocyte soma area and branch length for astrocytes in co-cultures treated with and without LPS. (F) Schematic representation of monocultures of astrocytes incubated with the extracellular conditioned medium of LPS-stimulated MGLs or respective controls, as well as astrocyte RNA isolation and qRT-PCR analysis. (G-H) qRT-PCR analyses of reactive astrocyte markers showing mRNA expression of TNFα and CXCL1 in astrocytes co-cultured with conditioned media from control MGLs or astrocytes co-cultured with conditioned media from LPS treated MGLs. (I) Schematic of hPSC-derived RGC, astrocyte and MGL triple culture control or stimulated with LPS. (J-K’) Representative images of RGCs (MAP2), astrocytes (GFAP) and MGLs (IBA1) grown in triple culture with and without LPS for 1 and 3 weeks. (L-N) Quantification of the number of RGCs per field of view in triple co-cultures control or stimulated with LPS for 1 or 3 weeks, showing a reduction in the number of RGCs when grown in cultures stimulated with LPS. (O-P) Representative images of RGCs (MAP2) in triple co-cultures control and stimulated with LPS. (Q-R) Quantification of RGC morphology including sholl analysis and neurite length, showing a reduction in RGC neurite complexity when in triple cultures stimulated with LPS. (S-T) Representative images of astrocytes (GFAP) in triple cultures control or stimulated with LPS. (U-V) Quantification of astrocyte branch length and number of branches showing a decrease in astrocyte branching and extension when grown in triple cultures stimulated with LPS compared to controls. (W-X) Representative images of MGLs (IBA1) in triple co-cultures control or stimulated with LPS. (Y-Z) Quantification of MGL morphology showing a reduction in microglial endpoints and increase in microglia circularity in MGLs grown in triple co-cultures stimulated with LPS compared to controls. Data represent mean values ± SEM. *p < 0.05, **p < 0.01 and ***p < 0.0001. Error bars represent SEM (*p<0.05, **p<0.01, ***p<0.001, ****P<0.0001, ns = not significant). Scale bar equals 100 μm in B and J, 50 μm in O, S, and W.

Next, we wanted to determine whether these presumptive microglia-activated astrocytes expressed additional features associated with astrocyte reactivity. Microglial activation was stimulated with LPS for 24 hours, and then microglia conditioned media was collected and filtered from both control and LPS-activated microglia. Microglia conditioned medium was then added to astrocytes and qRT-PCR analyses were performed to assess mRNA expression levels of the pro-inflammatory factors TNFα and CXCL10 (12, 37). These experiments allowed for the direct assessment of how astrocytes may be induced to a pro-inflammatory state while excluding potential confounding effects of residual microglia-derived factors (Figure 6F). Results demonstrated that when conditioned medium from LPS-activated microglia was added to astrocytes, a significant increase in the expression of activated astrocyte markers TNFα and CXCL10 was observed (Figure 6G-H), suggesting that astrocytes could further contribute to a pro-inflammatory environment in response to signals from pro-inflammatory microglia.

Building upon our findings that LPS-stimulated microglia confer degenerative phenotypes upon RGCs, and also induce a pro-inflammatory profile in astrocytes, we next examined a tri-culture model of RGCs, microglia, and astrocytes in which microglia were stimulated to a pro-inflammatory state to investigate how this inflammatory environment modulated RGCs in a combined context (Figure 6I). Sustained microglial activation was induced by treating tri-cultures with LPS every 48 hours, and RGC viability was assessed at 1 and 3 weeks (Figure 6J-N). At 1 week, LPS-treated triple cultures showed a significant reduction in RGC viability compared to untreated controls. Quantification of MAP2+ RGCs revealed a marked decrease in the number of RGCs per field of view, consistent with early neurotoxic effects of activated microglia. By 3 weeks, neurotoxic effects were more pronounced, with a further decrease in the number of MAP2+ RGCs per field of view. Of the RGCs that remained in these tri-cultures, they displayed a more retracted morphology with reduced outgrowth and branching complexity (Figure 6O-R). Additionally, astrocytes in triple culture demonstrated signs of activation including increased soma area and decreased individual branch length compared to controls (Figure 6S-V), while microglia exhibit a more round “amoeboid-like” morphology (Figure 6W-Z) in response to the LPS stimulation. These findings demonstrated that chronic microglial activation in the presence of astrocytes leads to progressive, time-dependent RGC degeneration, suggesting that the combined pro-inflammatory effects of astrocytes and microglia exacerbate inflammatory signaling and RGC neurotoxicity.

### Single Cell RNA-seq profiling of RGCs, microglia, and astrocytes to identify transcriptional correlates underlying neuroinflammatory signatures in tri-culture

To further investigate how the pro-inflammatory environment resulted in changes to all three cell types compared to those grown in more homeostatic control tri-cultures, we conducted single-cell RNA sequencing analyses on RGCs, astrocytes, and microglia between these two experimental conditions (Figure 7A-B). In these analyses, we observed numerous transcriptional changes in each cell type across the two experimental conditions. Interestingly, whereas prior experiments comparing tri-cultures to monocultures of each cell type demonstrated the largest number of differentially expressed genes (DEGs) in RGCs, in the context of a pro-inflammatory environment we observed the largest number of DEGs within microglia (Figure 7C). However, while many of these microglia DEGs were downregulated in the tri-culture environment, both RGCs and astrocytes exhibited larger numbers of DEGs that were upregulated in the tri-culture environment (Figure 7C).

**Figure 7.**
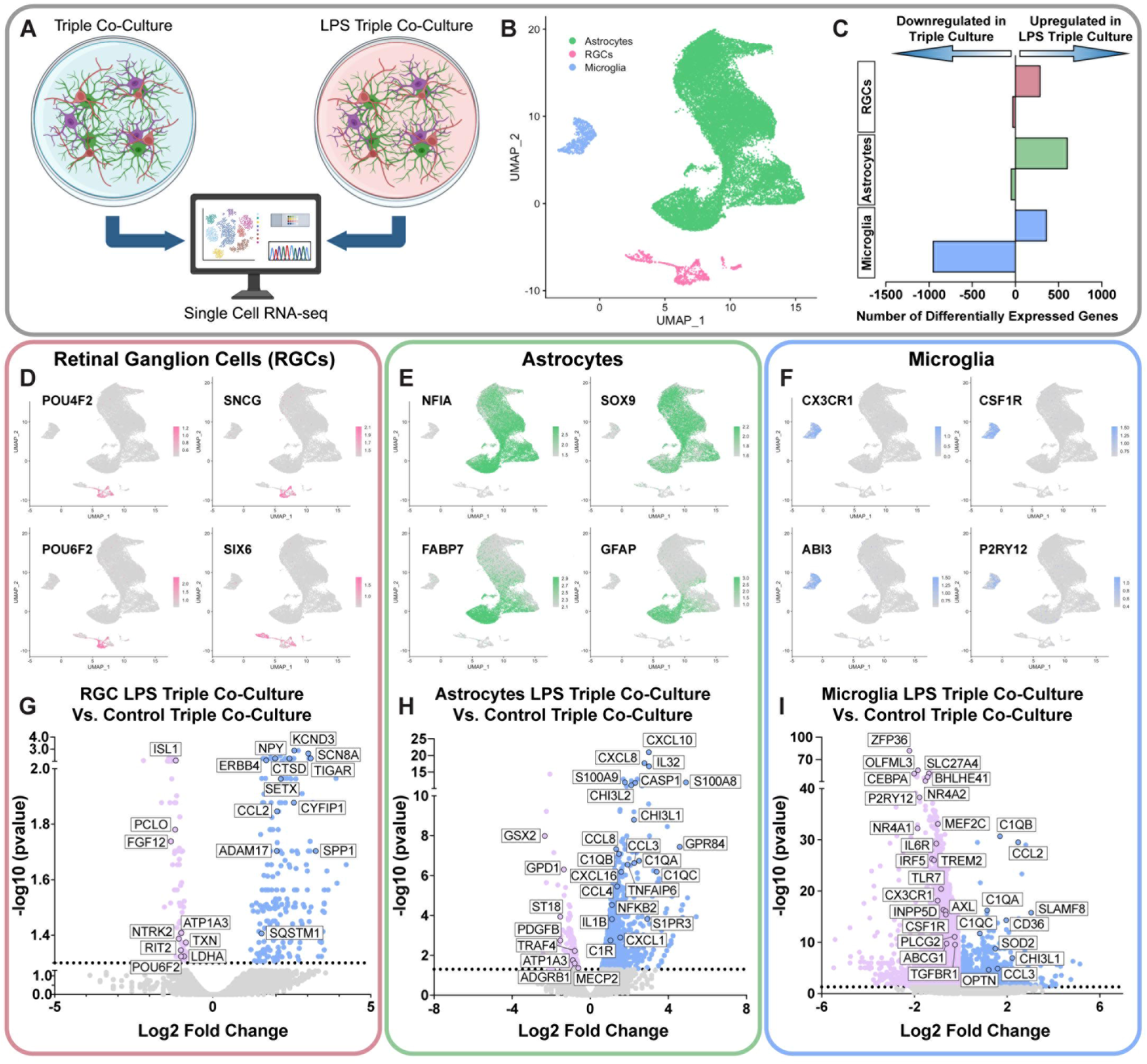
scRNA-seq of RGCs, astrocytes and MGLs to uncover transcriptional signatures associated with neuroinflammation in a tri-culture model. (A) Schematic representation of cell culture systems used for scRNA-seq including triple cultures containing RGCs, astrocytes and MGLs with or without treatment with LPS. (B) UMAP projection of scRNA-seq representing RGC, astrocyte and MGL population colored in pink, green and blue, respectively. (C) Representation of the number of differentially expressed genes triple-cultures untreated or treated with LPS. UMAPs represent (D) RGC populations (POU4F2, SNCG, POU6F2 and SIX6), (E) astrocyte populations (NFIA, SOX9, FABP7 and GFAP) and MGL populations (CX3CR1, CSF1R, ABI3 and P2RY12). Volcano plots exhibit differentially expressed genes in (G) RGC, (H) astrocytes, and (I) MGLs, when in triple co-cultures stimulated with LPS or control triple co-cultures. Significantly upregulated genes are shown in blue, while significantly downregulated genes are shown in pink.

Among RGCs in control tri-cultures compared to LPS-stimulated tri-cultures, we observed numerous transcripts upregulated in LPS-stimulated tri-cultures associated with neuroinflammation (CCL2, ADAM17, SPP1), cellular stress response (SQSTM1, CTSD, TIGAR), and neuronal excitability (KCND3, SCN8A, NPY). Conversely RGCs in control tri-cultures exhibited higher levels of transcripts associated with RGC survival (FGF12, NTRK2, TXN) and RGC identity (POU6F2, ISL1, RIT2), with the associated decrease in RGC identity genes in pro-inflammatory tri-cultures suggesting RGC degeneration (Figure 7G). Similarly, when comparing astrocyte populations, we found that astrocytes in the pro-inflammatory experimental group expressed higher levels of transcripts associated with a pro-inflammatory state (CXCL10, CCL3, CXCL1), complement pathway activation (C1QA, C1QC, C1R), and astrocyte reactivity (NFKB2, CHI3L2, CHI3L1), further validating the pro-inflammatory state of astrocytes conferred by pro-inflammatory microglia (Figure 7H). Finally, a comparison of microglia across experimental groups demonstrated that microglia in the pro-inflammatory condition expressed elevated levels of transcripts associated with a pro-inflammatory state (CCL2, CCL3, CHI3L1, SLAMF8) and complement pathway activation (C1QA, C1QB, C1QC), whereas control homeostatic conditions were associated with higher levels of transcripts in microglia associated with a homeostatic state (P2RY12, CX3CR1, INPP5D, CSF1R) and an anti-inflammatory phenotype (ZFP36, NR4A2, MEF2C, NR4A1) (Figure 7I). Collectively, these results demonstrated that together both microglia and astrocytes create a more potent pro-inflammatory environment that is highly neurotoxic to RGCs.

## DISCUSSION

Neuroinflammation in the CNS is a complex process which has been implicated in the pathogenesis of many neurodegenerative diseases, including glaucoma (38–40). In recent years, the neuroinflammatory responses associated with neurodegenerative disease have been shown to be mainly driven by reactive microglia and astrocytes (38). However, most studies on microglial activation have focused on animal models (30, 41–43), which do not fully recapitulate human disease. Some studies have also described morphological and functional differences in microglia, astrocytes and RGCs between rodent models and humans (44–48), including a variety of heterogeneous cellular phenotypes that can alter the function of these cell types depending upon their microenvironment. However, the complexity of the neuroinflammatory processes that may trigger or exacerbate RGC degeneration in glaucoma remains unclear, and we do not yet know if or how microglia and astrocytes contribute to the disease progression.

Microglial activation has been observed within the retina and optic nerve head (ONH) at the early stages of glaucoma, prior to RGC degeneration (30, 35). Previous studies have shown that treatment with the antibiotic minocycline (42) or irradiation (41) suppresses chronic neuroinflammation, reducing microglia activation and improving RGC pathology after injury. Furthermore, in an ocular hypertension model, the reduction of microglia activation by modulating microRNA-93-5p was neuroprotective to RGCs (49). Conversely, several other studies have demonstrated that PLX5622-induced microglial ablation exacerbates RGC loss following retinal or optic nerve damage, suggesting that microglia are neuroprotective (50, 51). Therefore, the contribution of microglia to RGC neurodegeneration in glaucoma remains controversial and the question of whether microglia are beneficial or harmful to RGCs is unknown. Thus, we developed a human cellular model of human RGCs and microglia (MGLs) to examine the effects of microglia activation on RGC morphology and function. Our results show that when RGCs are co-cultured for up to 3 weeks, microglia activation induces neurotoxic effects upon RGCs.

In glaucoma, microglial activation has been associated with changes in morphology, increased phagocytic activity, as well as the release of large amounts of inflammatory factors including TNF-α, IL-1β and IL-6, which promote the neurodegeneration of RGCs and tissue damage (30, 36, 41, 52, 53). As Toll-Like Receptor (TLR) signaling is one of the key inflammatory signaling pathways that has been shown to be upregulated in glaucomatous retinas (54), in this study we utilized a potent ligand (LPS) that initiates TLR-4 signaling (55) to transition microglia to a pro-inflammatory activated state. Here we show that once activated with LPS, hPSC-derived microglia (MGLs) alter their morphology, increase the expression of MHC-II, secrete various pro-inflammatory cytokines (TNFα, IL-1β, IL-6, IL-8, IFN-y) and enhance their phagocytic capacity when compared to control MGLs. Moreover, we observed that when analyzing the cytokines secreted from both control and LPS MGLs, some major anti-inflammatory cytokines (IL-4, IL-10 and IL-13) were also increased after LPS treatment. The increased secretion of both pro-inflammatory and anti-inflammatory cytokines is likely due to MGL heterogeneity *in vitro*, which could potentially be favorable when studying specific subsets of human microglia and how these heterogenous populations function in response to inflammatory triggers. Further, while LPS activation is often considered the gold standard to induce a microglial pro-inflammatory state, in future studies it will be important to test additional stimuli to yield a pro-inflammatory microglial phenotype, particularly those that may be more directly physiologically relevant than LPS.

Additionally, when assessing the phagocytic activity of control and LPS MGLs, we observed that LPS stimulation increased the ability of MGLs to phagocytose pHrodo Zymosan-A. Interestingly, microglia have previously been shown to be neuroprotective by increasing their phagocytic capacity to eliminate a variety of pathological material including pathogens, protein aggregates and cellular debris in order to maintain homeostasis (56). However, enhanced microglial phagocytosis has recently been shown to exacerbate neurodegeneration (57–59). Previous studies have shown that weak synapses of damaged neurons are tagged with complement components C1q and C3, which sends “eat me” signals to microglia initiating pathological pruning of stressed but viable neurons that may have had the potential to survive (60–62). Therefore, further studies are needed to understand the mechanisms by which increased microglial phagocytosis triggered by microglia activation could play a key role in the onset and progression of neurodegenerative diseases.

It has been shown that after treatment with LPS, microglia secrete 3 major inflammatory factors (IL-1α, TNF and C1q) that transition astrocytes toward a reactive state (12), which may be detrimental to neuronal survival and function. Our results show that when microglia activation is induced by LPS, astrocytes exhibit a neurotoxic reactive phenotype characterized by a reduction in the length of astrocyte branches, as well as an increase in the soma area, which is similar to what has been previously described for astrocytes that exhibit an “A1” phenotype. Furthermore, as reactive astrocytes have previously been shown to release inflammatory factors such as TNF-α and CXCL10 (12, 63), we observed that TNF-α and CXCL10 mRNA expression was also increased after treatment with LPS-MCM compared to control-MCM. Finally, when MGLs and astrocytes are cultured together with RGCs, both cells exert neurotoxic effects upon RGCs accelerating RGC neurodegenerative phenotypes when compared to LPS-activated microglia co-cultured with RGCs alone.

One limitation of our current model system is the use of a limited number of iPSC lines, which constrains our ability to fully capture the genetic and phenotypic diversity observed in the human population. Although findings were consistent across two independent lines, broader validation across additional genetic backgrounds will be necessary to assess the generalizability of these results. Moreover, while our tri-culture system successfully models key cellular interactions among RGCs, astrocytes, and microglia, it does not incorporate other components of the retinal environment such as vascular cells, systemic immune signals, or extracellular matrix cues that may influence glial activation and neuronal health i*n vivo*. The culture system also lacks the spatial architecture of the retina and optic nerve head, where glial reactivity is often regionally restricted. To better mimic the spatial orientation of activated glia along RGC axons akin to the optic nerve and/or optic nerve head, future studies will need to focus on utilizing alternative cell culture paradigms such as microfluidic platforms to better model the focal microglial activation observed along the proximal axonal compartment of RGCs, which is a defining feature of glaucomatous degeneration (35).

Additionally, while this tri-culture model was developed with a focus on the neuroinflammatory aspects of glaucoma, its relevance extends to a broader spectrum of neurodegenerative and neuroinflammatory disorders that affect the retina and optic nerve. Conditions such as optic neuritis, ischemic optic neuropathy, Leber hereditary optic neuropathy, and even retinal manifestations of systemic diseases like multiple sclerosis and Alzheimer’s disease involve glial activation and RGC degeneration. The ability to model human-specific interactions between microglia, astrocytes, and RGCs provides a powerful platform to dissect shared and disease-specific mechanisms of neuroinflammation and neuronal vulnerability. Moreover, because this system can be adapted using patient-derived iPSCs, it offers opportunities to study genetic forms of optic neuropathies or systemic disorders with retinal involvement, enabling the exploration of disease heterogeneity and the development of personalized therapeutic strategies.

In conclusion, our triple culture model will serve as a platform for the development of more precise human cellular models that can be used to study RGC/ glia interactions and their contributions to RGC degeneration. This human model system will also open new avenues in drug discovery, for the identification of therapeutic targets that could be used to treat retinal degenerative diseases such as glaucoma.

## MATERIALS AND METHODS

### Maintenance of hPSC cultures

hPSC lines used within this study included the H7 and WTC11 cell lines. For some experiments, fluorescent reporter cell lines were used (H7-Brn3b-tdTomato-Thy1). hPSCs were maintained on Matrigel or Geltrex-coated 6 well-plates in mTeSR1 or mTeSR+ medium to maintain pluripotency. hPSCs were passaged using ReLeSR every 6-7 days, at 60-70% confluency.

### Differentiation of hPSCs to microglia

Initially, hPSCs were differentiated into hematopoietic progenitor cells (HPCs) using the STEMdiff hematopoietic kit (StemCell Technologies), following previously established protocols (17, 64). Briefly, on Day-1, hPSCs were dissociated into small clusters of cells with ReLeSR into mTeSR1 or mTeSR1 PLUS medium. Clusters of cells approximately 100-200uM in diameter were counted in a 96 well plate and 40-60 clusters of cells were then plated onto reduced growth factor Matrigel-coated (1mg/mL) 6 well plates. On Day 0, colonies were counted and wells that had a density of 20-40 attached clusters were selected for HPC differentiation. mTeSR1 media was removed and differentiation was accomplished over the course of 10-12 days following manufacturer’s instructions. On Days 10 and 12, floating round, phase-bright HPCs were collected gently by collecting all medium from wells using a serological pipette. HPCs were then plated on reduced growth factor Matrigel coated 6-well plates at 80-100K cells per well in 2ml of microglia basal medium (DMEM/F12, 2X insulin-transferrin-selenite, 2X B27, 0.5X N2, 1X Glutamax, 1X non-essential amino acids, 400 μM monothioglycerol, 5 μg/mL insulin) supplemented with cytokines IL-34 (100ng/ml), TGFβ-1 (50ng/ml) and M-CSF (25ng/ml). 1ml of microglia basal medium was added with fresh cytokines every other day until Day 28. On Day 28, microglia-like-cells (MGLs) were harvested and used for experiments. To induce an activated phenotype, MGLs were incubated with 150 ng/mL of Lipopolysaccharide (LPS) (Millipore-Sigma) for 24 hours, based on previous protocols.

### Astrocyte differentiation from hPSCs

Astrocytes were differentiated from hPSCs following established protocols (21, 22, 65), which has been previously demonstrated by our group (21, 22, 65). Briefly, hPSC colonies were lifted using dispase to form free-floating cellular aggregates. On Days 0-3, cellular aggregates were slowly weaned off mTeSR1 medium and transitioned to neural induction media (NIM: DMEM/F12 with 1% N2 supplement and heparin, 2 ug/ml). On day 7, cellular aggregates were plated on a laminin-coated 6-well plate, with fresh NIM media changes every other day. On Day 16, neurospheres were carefully lifted from the plates and transferred to 10cm dishes in neurosphere differentiation medium (DMEM/F12, 2% B27 supplement, 1% MEM non-essential amino acids, and 1x anti-biotic, anti-mycotic). On day 40, neurospheres were transitioned to astrocyte differentiation medium containing fibroblast growth factor 2 (FGF2) (20 ng/mL) and epidermal growth factor (EGF) (20 ng/mL). We then matured astrospheres for a total of 6 months with mechanical passaging using a tissue chopper every 2 weeks. After 6 months, astrocytes were enzymatically isolated with Accutase to plate in 2D culture by dissociating the astrospheres into single cells and plating of cells on laminin and poly-ornithine coated 12mm coverslips, with 10,000 cells per coverslip. After plating, astrocytes were matured for an additional 3 weeks in serum-free BrainPhys culture medium containing 2% B27, 1% N2, ascorbic acid, cAMP, BDNF (20ng/ml), GDNF (20ng/ml) and CNTF (20ng/ml) (21, 22).

### Retinal ganglion cell differentiation

Retinal organoids were differentiated using our previously established retinal organoid protocol (19). Briefly, hPSCs colonies at approximately 70% confluency were dissociated into single cells using Accutase and seeded 2000 cells per well of a 96-well low adhesion U-bottom plate. Cells were seeded into wells in 100μL of mTeSR1 with 20μM ROCK inhibitor (Y-27632). The next day (Day 0) 50uL of mTeSR1 and 50uL of Neural Induction Medium (NIM) comprised of (DMEM/F12, 1% N2 supplement, 1% MEM non-essential amino acids, 1% anti-anti, and (2 μg/ml) heparin) with 20μM ROCK inhibitor (Y-27632) was added to each well. On Day 1 of differentiation cellular aggregates were removed from the 96-well plates using a multi-channel pipette and transferred to a 10cm dish then gradually transitioned to complete NIM medium by Day 3. On Day 6, fresh NIM with 50ng/mL of BMP4 was added to the cellular aggregates to induce retinal specification. On Day 8 of differentiation, aggregates were plated on 6-well plates with the addition of 10% Fetal Bovine Serum. Half of the media was removed on Day 9, 12 and 15 and replaced with fresh NIM. On Day 16, retinal organoids were carefully removed from the plates and transferred to 10cm dishes containing Retinal Differentiation Medium (DMEM/F12, 2% B27 supplement, 1% MEM non-essential amino acids, and 1x anti-biotic, anti-mycotic) with 1% FBS. On day 18 and day 20, media was changed and RDM was supplemented with 3% FBS and 5% FBS respectively. On day 22, organoids were then transitioned to Advanced-RDM (ARDM) medium consisting of (DMEM/F12, 2% B27 supplement, 1% MEM non-essential amino acids, 1x anti-biotic, anti-mycotic, 1x GlutaMAX, 10% FBS and 100μM taurine. Organoid media was changed every 2-3 days. GlutaMAX, FBS and Taurine were added fresh each media change. At Day 40-45, retinal organoids were enzymatically dissociated and RGCs were purified from the organoids using accumax for 30 minutes at 37°C and using Magnetic Activated Cell Sorting (MACS) and the Thy1.2 cell surface antigen (Miltenyi), RGCs were purified and plated on laminin and poly-ornithine coverslips at 10,000 cells per coverslips. After plating, RGCs were maintained in serum-free BrainPhys culture medium containing 2% B27, 1% N2, ascorbic acid, cAMP, BDNF (20ng/ml), GDNF (20ng/ml) (18, 66).

### qRT-PCR analyses

For qRT-PCR analyses, RNA was collected for iPSCs, astrocytes, cortical neurons or MGLs. Total RNA was extracted using the Picopure RNA extraction kit (Life Technologies) following manufacturer’s instructions. cDNA was then synthesized using the High-Capacity RNA-to-cDNA Kit (Applied Biosystems) and qRT-PCR analyses was performed using a Quant Studio 7 Flex system (Applied Biosystems) with SYBR green (Life Technologies), using primers as specified in Supplemental Table 1. Gene expression was defined using a comparative Ct method, normalized to β-actin. The ΔCt value was normalized to the average ΔCt value of the traditional method day 30 organoids to calculate ΔΔCt and fold change was calculated using the formula (2^−ΔΔCt^), as previously described (67).

### Co-cultures of hPSC-derived RGCs, Microglia and Astrocytes

Co-cultures were established between RGCs and microglia at a 5:1 ratio, while triple cultures were established between RGCs, microglia and astrocytes at a 5:1:2 ratio. These co-cultures were maintained in serum-free BrainPhys culture medium containing 2% B27, 1% N2, Glutamax (1X) and BDNF (20ng/ml), GDNF (20ng/ml) and CNTF (20ng/ml) (Miltenyi) (18, 66) with the addition of cytokines IL-34 (100ng/ml), TGFβ-1 (50ng/ml) and M-CSF (25ng/ml). To induce microglia activation in co-cultures, cultures were incubated with 150 ng/mL of Lipopolysaccharide (LPS) every 48 hours and TGFβ-1 was excluded from the media.

### Immunocytochemistry and Imaging

Microglia differentiation was determined to be successful using brightfield imaging, as well as using immunostaining for P2RY12, IBA1 and TREM2. Microglia were fixed for 15 minutes in 4% paraformaldehyde, then washed 3x with 1xPBS for 5 minutes each. Immunostaining of MGLs was then performed as previously described. Primary antibodies used are listed in Supplemental Table 2. Samples were imaged either on a Leica DM5500 upright microscope, or a Nikon AR1 confocal microscope or confocal.

### MesoScale Discovery (MSD) multiplex ELISA analyses

To analyze the secretion of inflammatory factors released from both control and activated MGLs, microglia were differentiated until Day 28 from HPCs, then incubated 24 hours with LPS (150ng/ml) or maintained as controls. We then collected conditioned media for both control and activated MGLs and filtered the media with a 0.2 µm filter before analyzing the conditioned medium from both populations using the Mesoscale Discovery V-Plex proinflammatory panel, as previously described.

### Phagocytosis assays

To determine changes in the ability of activated MGLs to phagocytose material, both control and activated MGLs were incubated with pHrodo Red Zymosan BioParticles at a final concentration of 50 ng/ml. Cells were then imaged using time-lapse imaging on the Lux microscope and imaged every 5 minutes for 12 hours. The Lux software quantified the red fluorescence overtime (red object count) which was then normalized to the total number of cells per field of view.

### Quantification and statistical analysis

Statistical significance of results was determined using either a Student’s t-test or a one-way ANOVA followed by Tukey’s post-hoc analysis based on a p-value < 0.05 using Prism software v. 9.5.1. A minimum of 3 biological replicates were used for all experiments. The circularity of MGLs was quantified using Image J by calculating the area (mm^2^) and circumference. Circularity was quantified as a measure of cell shape using the formula: Circularity = Perimeter^2^/4 × π × Area (4*(3.14)*(Area)/(Perimeter^2). RGC and Astrocyte morphologies were quantified as previously described (20).

Microglial morphology was analyzed using the AnalyzeSkeleton (2D/3D) plugin in ImageJ (v1.54g, NIH, USA), following the protocol described by Young & Morrison (68). A minimum of seven images per condition were analyzed to measure average process length and the number of endpoints per microglial cell.

### Preparation of Tri-Cultures for Single Cell RNA-sequencing

Astrocytes were plated into 12 well laminin coated plates and matured for 2 weeks in serum-free BrainPhys culture medium containing 2% B27, 1% N2, Glutamax (1X) and BDNF (20ng/ml), GDNF (20ng/ml) and CNTF (20ng/ml) (Miltenyi) (18, 66). After 2 weeks of maturation, RGCs were added to astrocyte cultures and both RGCs and astrocytes were further matured for an additional week. After 3 weeks, MGLs were added, and triple cultures were maintained for 1 week total prior to performing single cell RNA-sequencing. To induce microglia activation in triple cultures, LPS (150 ng/mL) was added every 48 hours.

Single cells in suspension were prepared from 1 week triple cultures with the addition of 1mL of accutase per well. Cells were incubated with accutase for 15–20 min at 37^ο^C to ensure dissociation of all cells. After complete dissociation, accutase was diluted in 5ml of Brain Phys medium containing BDNF, GDNF, CNTF, IL-34 and mCSF + 10uM ROCK inhibitor and the cell suspension was run through a 30-μm strainer to remove clumps. The cell suspension was then spun down at 300xg for 5 minutes and the cell pellet was then resuspended in 70 μL of Brain Phys medium containing BDNF, GDNF, CNTF, IL-34 and mCSF + 10uM ROCK inhibitor, placed on ice, and transferred to the core facility for further processing. The cell suspension was washed at least 3 times with PBS plus 0.04% BSA and cell viability was then assessed to a targeted cell concentration of 700– 1200 cells/μL, as recommended in the 10x Genomics guidelines.

The single cell analysis was conducted using a 10X Chromium single cell system (10X Genomics, Inc) and a NovaSeq 6000 sequencer (Illumina, Inc). Following the CG000315_ChromiumNextGEMSingleCell3 GeneExpression_v3.1_DualIndex (10X Genomics, Inc), cell suspension along with RT reagents, and the single cell gel beads and partition oil, were transferred onto a Next GEM Chip G in separate wells, one sample per well, and the chip loaded to the 10X Controller for GEM generation and barcoding, followed by RT reaction. cDNA synthesis and cDNA library preparation. At each step, the quality of cDNA and cDNA library were examined by Bioanalyzer and Qubit. The resulting cDNA libraries were sequenced on an Illumina NovaSeq 6000, with paired-end 100 bp generated. Depending on targeted cell recovery, about 50K paired reads per cell were generated and 93% of the sequencing reads reached Q30 (99.9% base call accuracy). A Phred quality score (Q score) was used to measure the quality of sequencing.

The sequence data were first processed with cellranger (7.2.0). The human reference genome GRCh38-2020-A was used for the sequence data alignment. The feature-cell barcode matrices generated were used for further analysis.

### Single Cell RNA-sequencing Analysis

Single-cell RNA sequencing (scRNA-seq) libraries were generated and sequenced on an Illumina platform. The resulting.h5 gene expression matrices which contained the raw gene by cell count data were provided by a genomics core sequencing facility. For initial quality control, h5 files were uploaded to Trail Maker, the Parse Biosciences Cloud Analysis Platform which supports the preprocessing and quality control of scRNA-seq data. Quality control and principal component analysis was performed using the Trail Maker Pipeline. QC thresholds are automatically computed per sample, based on global and per-sample distributions of gene count, transcript count, and mitochondrial fraction to ensure consistent retention of high-quality cells. The.rds file containing a preprocessed Seurat object, which includes raw count matrix, normalized expression data, metadata, and dimensionality reduction embeddings were then imported in R (v4.4.2 or later) for downstream analysis using the Seurat package. Differential gene expression (DGE) analysis was performed to identify cluster-specific transcriptional signatures within specific cell types (RGCs, Astrocytes, and Microglia). After normalization, dimensionality reduction, and clustering, major cell types were identified using canonical marker genes and assigned to each cluster. To compare treatment conditions (control vs. LPS) within each annotated cell type, cells were subset by cluster identity and treatment metadata. DEG analysis was performed using Seurat’s FindMarkers() function, which applies the Wilcoxon rank-sum test. The comparison was conducted within each cell type, using the group.by and subset.ident arguments to isolate the relevant subset of cells. Results were filtered based on log2FC and p-values (p_val < 0.05) and interpreted to identify genes significantly affected by the treatment in a cell-type-specific manner.

## ACKNOWLEDGMENTS

We thank Hongyu Gao and Fang Fang at the Indiana University Center for Medical Genomics for assistance with single cell RNA-seq experiments, as well as Akshayalashmi Sridhar and Subramanian Dharmarajan for assistance with single cell RNA-seq analyses. Grant support was provided by the National Eye Institute (R01EY033022 and U24EY033269 to JSM), the Gilbert Family Foundation (923016 to JSM), and the BrightFocus Foundation (G2020369 to JSM). Support for this project was also provided by the Sarah Roush Memorial Fellowship from the Indiana Alzheimer’s Disease Center (CG) and the BrightFocus Postdoctoral Fellowship (G2022003F to CG), as well as a Cagiantas scholarship from the Indiana University School of Medicine (JH).

## Notes

### Competing Interest Statement

The authors have declared no competing interest.

