## Supplemental Information for "A human pluripotent stem cell tri-culture platform to elucidate microglial regulation of retinal ganglion cells in neuroinflammation"

**Supplemental Table 1.** List of qRT-PCR primers used in the study.

| **Gene Amplified** | **Forward sequence** | **Reverse sequence** |
| --- | --- | --- |
| β-Actin | GCG AGA AGA TGA CCC AGA TC | CCA GTG GTA CGG CCA GAG G |
| P2RY12 | CCT GGA TCC GTT CAT CTA TTT | CAC CAC CAT CCT GTT CTT T |
| TREM2 | GGC TGC TCA TCT TAC TCT TT | GTG CTT CAT GGA GTC ATA GG |
| IBA1 | TGT CTC CCC ACC TCT ACC AG | CCT TCA GCA GTC CGA AAG CT |
| PU.1 | CAC GGA TCT ATA CCA ACG CCA | CCA GTA ATG GTC GCT ATG GC |
| CSFR1 | CCA GGC TAA AAG GGG AAG AAG | CGT TCC TCT CCT CTG CAC T |
| TNF-a | CAC AGT GAA GTG CTG GCA AC | AGG AAG GCC TAA GGT CCA CT |
| CXCL10 | TGT ACG CTG TAC CTG CAT CA | ACG TGG ACA AAA TTG GCT TGC |
| CD74 | GAT GAC CAG CGC GAC CTT ATC | GTG ACT GTC AGT TTG TCC AGC |
| ABI3 | ATC GCC CCA GAG AAC CTA CC | GCT CTT TCG AGA CAG GGT GC |

**Supplemental Table 2.** List of antibodies used in the study.

| **Antibody** | **Type** | **Source** | **Catalog** | **Dilution** |
| --- | --- | --- | --- | --- |
| GFAP | Chicken polyclonal | Aves labs | GFAP87987979 | 1:100 |
| IBA1 | Rabbit polyclonal | Fijufilm Wako Chemicals | 019-19741 | 1:500 |
| IBA1 | Goat polyclonal | Abcam | Ab5076 | 1:100 |
| MAP2 | Mouse monoclonal | Synaptic Systems | 188011 | 1:200 |
| P2RY12 | Rabbit polyclonal | Sigma Aldrich | HPA014518 | 1:200 |
| TREM2 | Goat polyclonal | R&D Systems | AF1828 | 1:200 |
